# Simple, rapid, and sensitive quantification of dolichyl phosphates using phosphate methylation and reverse-phase liquid chromatography-high resolution mass spectrometry

**DOI:** 10.1101/2022.09.14.504994

**Authors:** Dipali Kale, Frauke Kikul, Prasad Phapale, Lars Beedgen, Christian Thiel, Britta Brügger

## Abstract

Dolichyl monophosphates (DolPs) are essential lipids in glycosylation pathways that are highly conserved across almost all domains of life. The availability of DolP is crucial for all glycosylation processes, as these lipids serve as membrane-anchored building blocks that various types of glycosyltransferases use to generate complex post-translational modifications of proteins and lipids. Analysis of DolP species by reverse-phase liquid chromatography-mass spectrometry (RPLC-MS) has remained challenging due to their very low abundance and wide range of lipophilicities. Until now, a method for the simultaneous qualitative and quantitative assessment of DolP species from biological membranes has been lacking. Here we describe a novel approach based on simple sample preparation, rapid and efficient trimethylsilyl diazomethane (TMSD)-dependent phosphate methylation and RPLC-MS analysis for quantification of DolP species with different isoprene chain lengths. We used this workflow to selectively quantify DolP species from lipid extracts derived of *Saccharomyces cerevisiae*, HeLa and human skin fibroblasts from steroid 5-*α*-reductase 3-congenital disorders of glycosylation (SRD5A3-CDG) patients and healthy controls. Integration of this workflow with global lipidomics analyses will be a powerful tool to further our understanding of the role of DolPs in pathophysiological alterations of metabolic pathways downstream of HMG-CoA reductase, associated with CDGs, hypercholesterolemia, neurodegeneration, and cancer.

## Introduction

DolPs are long carbon chain lipids composed of alpha-saturated polyisoprenoid phosphates. DolPs with 18 - 21 isoprene units have been described in mammals [1], and are present in almost all eukaryotic membranes [2]. DolP species play crucial roles as glycan carriers in cellular glycosylation reactions where they serve as membrane-anchored acceptors for saccharides, presenting sugar substrates to a broad set of different transferases and allowing the formation of complex glycan structures [2]. Genetic defects in glycosylation constitute a large group of >165 diseases, referred to as CDGs, and result in a broad range of metabolic dysfunctions associated with defects in the process of carbohydrate addition to proteins or lipids [3]. Genetic mutations affecting DolP synthesis and recycling result in DHDDS-, NUS1-, SRD5A3-, and DOLK-CDGs, classified in the CDG-I subgroup of diseases [4–7]. Altered endogenous DolP levels have also been reported in several other pathological and physiological conditions, including Alzheimer’s disease, dementia and Prion disease [8–11]. DolP accounts for up to 20% of the total cellular dolichol pool [12, 13], and undergoes rapid metabolism as part of the dolichol cycle. *De novo* synthesis of dolichol via isoprenoid precursors in eukaryotes, archaea and some prokaryotes, is initiated by the mevalonate pathway, which is also utilized for the synthesis of cholesterol, heme and ubiquinone (Fig. S1). Prenylation of farnesyl diphosphate represents the committed step towards the synthesis of polyprenol, which is reduced to dolichol by steroid 5-*α*-reductase 3 (SRD5A3). DolK then converts dolichol to DolP, which in turn serves as an acceptor for monosaccharides, including mannose. DolP-mannose (DolP-Man), synthesized by the DolP-Man synthase, forms a critical metabolic hub that channels the mannose unit into the major glycosylation pathways localized at the endoplasmic reticulum, i.e. N-glycosylation, O- and C-mannosylation [14, 15] and glycosylphosphatidylinositol anchor synthesis [16]. Both *de novo* synthesis of DolP and its recycling following the transfer of the monosaccharide or glycan structures to acceptor proteins or lipids can adapt steady-state levels of DolP to the cellular demand. DolP with 19 isoprene units has been shown to orientate perpendicularly to the plane of the lipid bilayer, with the phosphate headgroup being facing the hydrophilic interface [17, 18]. The intercalation of DolP was shown to perturb the fatty acyl chain order of phosphoglycerolipids [17, 18]. It has been suggested that these biophysical properties of long-chain DolP species might facilitate membrane remodeling events that occur during the translocation of lipid-bound glycans across membranes and during membrane fusion [18, 19]. In addition, both dolichol and DolP have been reported to accumulate in membranes during aging [20], and DolP has been demonstrated to be a rate-limiting metabolite for cellular glycosylation and thereby essential for cell proliferation and development [21]. The accurate identification and quantification of DolPs at the cellular level using powerful analytical techniques such as MS is therefore crucial to understand mechanisms implicated in DolP metabolism and glycosylation pathways.

Conventionally, two separate analytical platforms have been used for cellular DolP analysis, firstly normal-phase liquid chromatography (NPLC) combined with fluorescence detection [1, 22] for quantification, and secondly MS-based workflows (see for example [22–26]) for structural characterizations. A frequently used, robust quantification method for DolPs by Hennet et al. [22] is based on the conversion of nonfluorescent DolPs to their fluorescent derivatives in a multi-step procedure, which is a labor intensive and time-consuming process for routine analyses. Methods based on MS workflows [22–26] mostly contain qualitative analyses of DolPs, which are often poorly ionized and suppressed by highly abundant lipid species [27, 28]. To our knowledge, there are currently no available methods for the simultaneous qualitative and quantitative assessment of DolP species from biological membranes.

RPLC-based approaches are widely used in metabolomics and lipidomics platforms, owing to their versatility, robustness, and MS compatibility [29–32]. (RPLC-)MS analysis of DolP species are, however, complicated by the following factors: (1) sensitivity; DolPs represent only a small percentage (~0.1% in eukaryotes) of total cellular phospholipids and thus have a very low abundance; (2) chemical diversity; DolP species cover a wide and high range in lipophilicity with an octanol:water partition coefficient (LogP) value distribution over 14 units (from 22.4 of DolP C65 to 37.4 of DolP C105); (3) dynamic range; individual DolP concentrations vary in human cells from 10.8 μg/g in the liver to 169 μg/g in testis [33].

We addressed these challenges by combining TMSD-based phosphate head group methylation with RPLC-MS analysis to simultaneously characterize and quantify DolPs from biological samples. TMSD methylation has been demonstrated to significantly increase the sensitivity of detecting different phospholipids, including low abundant phosphoinositides or bis(monoacylglycerol-) phosphates, which overcome the challenges mentioned earlier [34–41]. Here, we describe for the first time that TMDS methylation of DolPs can significantly improve the sensitivity of quantification through enhanced MS ionization efficiency and specificity by enabling their RPLC separation from co-eluting peaks. The developed RPLC-MS method provides high sensitivity (LOD ~1 pg), a broad molecular coverage, and a wide dynamic range. In addition, the overall sample preparation workflow used in this study is simple, fast, cost- and labor-effective, and does not require further steps of DolP enrichment such as solid phase extraction columns. As a proof of principle, we have applied this workflow to profile DolP from HeLa cells and *S. cerevisiae*, and to determine DolP levels in fibroblasts from SRD5A3-CDG patients and controls.

## Experimental section

### Reagents, chemicals, and lipid standards

DolP mixture (#900201X, hereinafter referred to as DolP standard) was purchased from Sigma-Aldrich (Merck KGaA, Darmstadt, Germany). Polyprenyl monophosphate (PolP) C60 standard was obtained from CymitQuimica (#48-62-1060, Barcelona, Spain). Potassium hydroxide (KOH), dichloromethane, and TMSD (#362832, 2 M in hexane) were obtained from Sigma-Aldrich. LC-MS grade water, isopropanol, methanol (MeOH), acetonitrile, ammonium acetate, and formic acid were from Fisher Scientific.

### Cell culture

HeLa cells were cultured in alphaMEM medium (Sigma Aldrich) supplemented with 10% fetal calf serum (FCS, Bioconcept, Switzerland), 2 mM glutamine, and penicillin-streptomycin (#P4333, Sigma Aldrich) and incubated in a humidified 5% CO_2_ atmosphere at 37 °C. For *S. cerevisiae* (strain W303-1B, derived from strain W303 [42]), approximately 5×10^7^ cells were cultured in YPD medium and harvested from cultures grown to mid-log phase with an OD_600_ value of 0.8. Fibroblasts from human patients were obtained with the patients’ parents written informed consent. This study was performed in accordance with the Declaration of Helsinki and approved by the Ethics Committee of the Medical Faculty Heidelberg (S-904/2019). The cells were maintained at 37 °C in a humidified atmosphere under 5% CO_2_. Patient and control fibroblasts were cultured in Dulbecco’s modified Eagle’s medium (high glucose; Life Technologies) supplemented with 10% FCS (PAN Biotech, Aidenbach, Germany), 100 U/mL penicillin, and 100 μg/mL streptomycin. Cell monolayers with 80 to 90% confluence in a T75 flask were washed twice with PBS and harvested using 2 mL 0.25% (v/v) trypsin-EDTA solution. All cell pellets were shock-frozen in liquid nitrogen and stored at −80 °C until lipid extractions were performed.

### Alkaline hydrolysis and lipid extraction

For the analysis of DolP from HeLa, *S. cerevisiae*, and human fibroblasts, the sample preparation and extraction protocol by Haeuptle et al. [22] was used with few modifications. Briefly, ~ 1×10^6^ HeLa or fibroblast cells were collected by trypsinization and washed with the medium (HeLa) or PBS (fibroblasts). *S. cerevisiae* cells were resuspended in 155 mM ammonium bicarbonate and homogenized using beads (0.5 mm; BioSpec Products). Lysates were diluted to 0.8 OD units (~ 8×10^6^ cells) per 200 μL. DolP extractions were performed in 13×100 mm Pyrex glass tubes. Twenty pmol PolP C60 was added to cell pellets as internal standard (IS), followed by 1 mL MeOH and then 1 mL H_2_O. Phosphate esters were hydrolyzed by adding 0.5 mL 15 M KOH, followed by an incubation of the samples for 60 min at 85 °C. Phase partitioning was induced by adding 1 mL MeOH and 4 mL dichloromethane, and lipids were further hydrolyzed for 60 min at 40 °C. The lower phase was washed four times with 2.7 mL dichloromethane/MeOH/H_2_O (3:48:47, v/v/v) and evaporated to dryness in a Turbovap evaporator chamber. In order to normalize sample amounts, phosphatidylcholine (PC) as bulk membrane lipid was extracted and analyzed as described [43]. Subsequent MS analysis was typically performed immediately after extraction. If measurements had to be postponed, lipid extracts were stored as lipid films under nitrogen at −20°C for a maximum of 3 days.

### Methylation of DolPs

For methylation of DolP in the above lipid extracts or standards, the TMSD reagent was used with necessary precautions. Dried samples were dissolved in 200 μL dichloromethane: MeOH (6.5:5.2, v/v). Then, 10 μL of TMSD was added, and samples were incubated for 40 min at RT. Following this incubation, 1 μL acetic acid was added to neutralize an excess of TMSD [44]. For RPLC-MS measurements, dried samples were reconstituted in 100 μL MeOH.

#### Safety considerations for handing TMSD

TMSD must be handled with care, as its inhalation may cause lung damage or central nervous system depression. It was used with the appropriate safety procedures and personal safety protective equipment inside the fume hood. Excess TMSD in the hood atmosphere was neutralized with acetic acid.

### Nano-electrospray direct infusion MS (nESI-DIMS) analysis

DolP standard and PolP C60 were methylated separately, evaporated to dryness, and resuspended in 60 μL each of 50 mM ammonium acetate in MeOH. As a control, the unmethylated DolP standard was prepared in the same way but omitting the TMSD methylation step. Both methylated and un-methylated DolPs (each at a final concentration of 42 ng/ μL) were then mixed with methylated PolP C60 (6 pmol/μl) in 25 mM ammonium acetate in chloroform: MeOH (1:1 v/v), transferred to a 96-well plate (Eppendorf, Hamburg, Germany). Five μL of the above sample was infused by nESI direct infusion using a TriVersa NanoMate (Advion Biosciences, Ithaca, NY, USA) coupled to a Q-Exactive Plus high-resolution MS (Thermo Scientific, MA, USA) (nESI-DIMS) for 7 min. The positive ionization voltage was set to 1.75 kV, and back pressure was 0.65 psi. MS analysis was performed in a range of m/z 930-1579 using a maximal injection time (max IT) of 3000 ms, automatic gain control for an ion target (AGC target) of 5×10^6^, resolution setting at 200 m/z (Rm/z200) = 1.4×10^5^ and ion transfer capillary temperature was set to 200 °C. Data analysis was performed using LipidXplorer [45], and internal standard-based normalization of peak intensities of DolPs was performed. The methylation efficiency was calculated using the following equation.

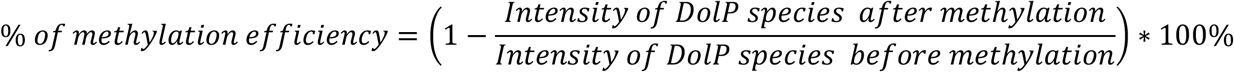

### RPLC-MS analysis

Separation of methylated DolPs was performed on a Dionex Ultimate 3000 UHPLC system coupled to a Q-Exactive Plus HRMS (RPLC-MS) operating in positive ionization mode. RPLC separation of methylated DolPs was carried out on a Waters CSH C18 column (1×150 mm, 1.7 μm) at a 0.1 mL/min flow rate, maintained at 55 °C. The mobile phase consisted of solvent A (acetonitrile:water (6:4, v/v)) and solvent B (isopropyl alcohol:acetonitrile (9:1, v/v)), buffered with 10 mM ammonium acetate and 0.1% formic acid. The following gradient profile was applied: 40% B at 0 min, 50% B in 3 min, 54% B in 9 min, 70% B in 9.1 min, 90% B in 17 min, 90% B for 27.5 min, 40% B in 27.6 min, and then 40% B was maintained up to 30 min. Lipids were detected with HRMS full scan MS (AGC target of 3×10^6^, Rm/z200 = 7×10^4^) and with parallel reaction monitoring (PRM) mode (max IT= 100 ms, AGC target of 2×10^5^, Rm/z200 = 1.75×10^4^, isolation window 4.0 m/z, and NCE 50) of 10 lipids at selected m/z mentioned in Table 1. The following MS parameters were set: Spray voltage on a HESI II ion source of 3.5 kV, sheath gas 30, auxiliary gas 10, spare gas 1 unit, S-Lens 50 eV, and capillary temperature 250 °C. The data analysis and processing were performed by Thermo Qual/Quan Browser Xcalibur (v 3.1; Thermo Fisher Scientific, Waltham, MA, USA). Peak areas of monoisotopic peaks were corrected for natural ^13^C isotope abundance. Relative peak area ratios of DolPs to IS were converted to molar concentrations using a known IS concentration. Statistical analysis of datasets was carried out using Microsoft Excel 2016 to perform Student t-tests. All graphs were generated using GraphPad Prism version 5.0.3 (GraphPad Software, Inc.).

**Table 1.** DolP and PolP molecular compositions and positive ion mode RPLC-MS characteristics of annotated di-methylated analyte species in this study.

| Analyte species | Isoprene units (n) | Molecular formula | Theoretical $m/z$ | Observed $m/z$ | Mass error (ppm) | RT (min) | RRT<sup>a</sup> |
| --- | --- | --- | --- | --- | --- | --- | --- |
| PoIP C60 | 12 | C62H103O4P1 | 960.7932 | 960.7924 | -0.8 | 19.1 | 1.00 |
| DoIP C65 | 13 | C67H113O4P1 | 1030.8715 | 1030.8711 | -0.4 | 20.0 | 1.05 |
| DoIP C70 | 14 | C72H121O4P1 | 1098.9341 | 1098.9360 | 1.7 | 20.6 | 1.08 |
| DoIP C75 | 15 | C77H129O4P1 | 1166.9967 | 1166.9976 | 0.8 | 21.2 | 1.11 |
| DoIP C80 | 16 | C82H137O4P1 | 1235.0593 | 1235.0613 | 1.6 | 21.8 | 1.14 |
| DoIP C85 | 17 | C87H145O4P1 | 1303.1219 | 1303.1229 | 0.8 | 22.3 | 1.17 |
| DoIP C90 | 18 | C92H153O4P1 | 1371.1845 | 1371.1854 | 0.7 | 22.9 | 1.2 |
| DoIP C95 | 19 | C97H161O4P1 | 1439.2471 | 1439.2476 | 0.3 | 23.5 | 1.23 |
| DoIP C100 | 20 | C102H169O4P1 | 1507.3097 | 1507.3127 | 2.0 | 24.1 | 1.26 |
| DoIP C105 | 21 | C107H177O4P1 | 1575.3720 | 1575.3724 | 0.3 | 24.7 | 1.29 |

<sup>a</sup> RRT, ratio of the retention of the peak relative to PoIP C60

## Results and discussion

### Increased sensitivity of DolPs detection through methylation

TMSD methylation has been shown to be a powerful approach to enhance the ionization efficiency of anionic phospholipids in the positive ion mode by eliminating the negative charge of the phosphate group [34, 36]. To test whether methylation of DolP species increases the sensitivity of MS detection, we used a commercial DolP mixture containing chain lengths from C65-C105 as a reference standard. Since the unmethylated DolP standard cannot be analysed by RPLC-MS, we used direct infusion nESI-DIMS analysis of both un-methylated and methylated DolPs to evaluate their ionization and methylation efficiency. As observed for anionic phospholipids, nESI DIMS analyses showed that un-methylated ions were poorly ionized, whereas methylation of DolP species resulted in a marked increase in ion intensities compared to the un-methylated standard (Fig. S2). However, with nESI-DIMS, we could only detect and quantify four abundant DolP species C80-C95, other species, although present in the DolP standard, were not detected. nESI-DIMS analysis of the DolP standard before TMSD methylation showed four [M+NH_4_]^+^ ions, i.e. DolP C80 at m/z 1207.0309, DolP C85 at m/z 1275.0907, DolP C90 at m/z 1343.1530, and DolP C95 at m/z 1411.2139 (Fig. 1A). After methylation, the ion cluster was shifted to m/z 1235.0630, 1303.1233, 1371.1881, and 1439.2474, respectively (Fig. 1B). The mass addition of ~ 28.0333 Da indicates di-methylation (addition of C2H4; exact mass = 28.0313 Da) of the phosphate residue in the DolP species. After methylation, un-methylated or mono-methylated species were not detected (see also Fig. S2, theoretical m/z values of un-, mono-, di-methylated DolP ions are given in Table S1).

**Figure 1.**
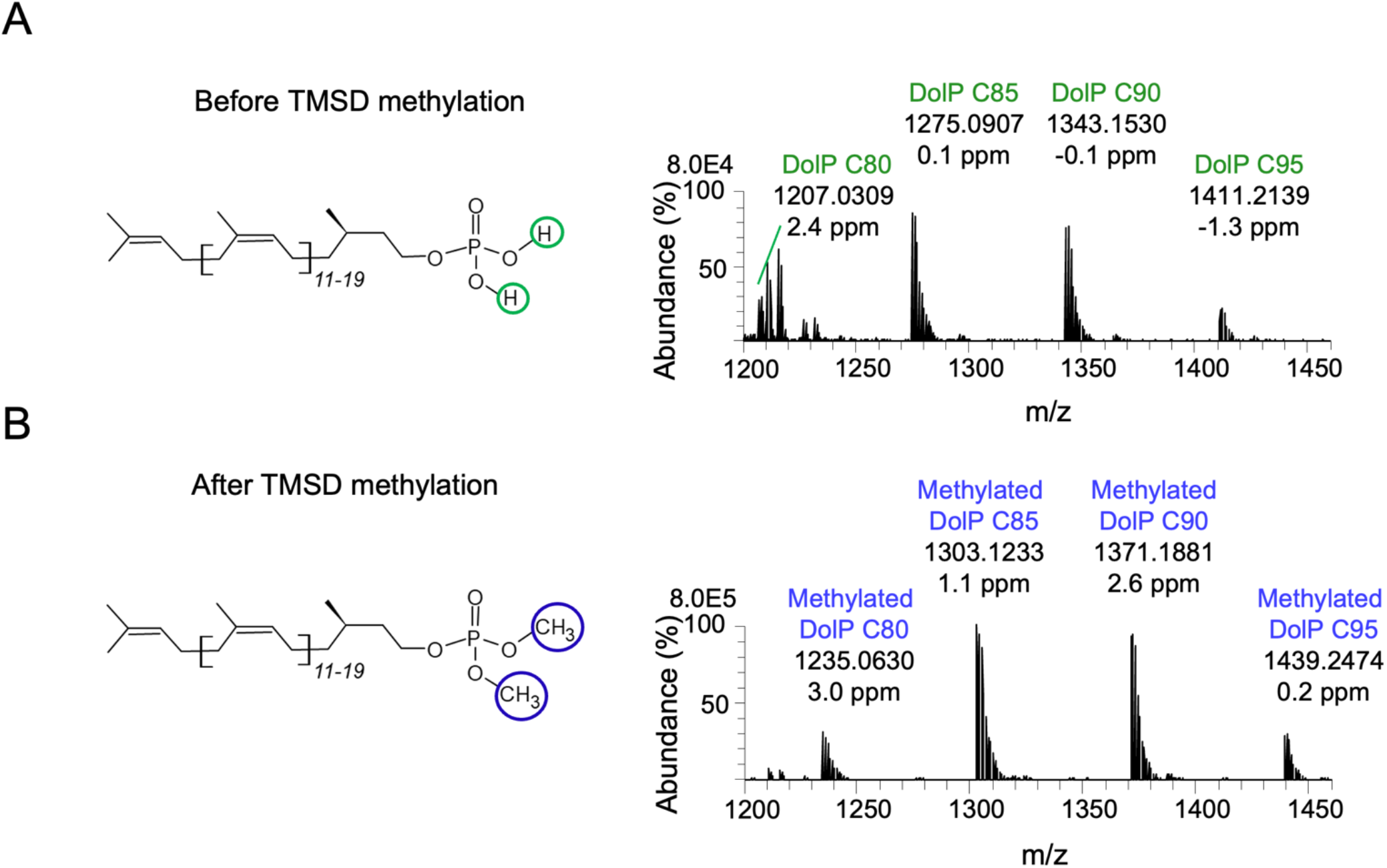
Representative positive ion nESI-DIMS spectra of DolP standard before and after methylation. Un-methylated (A) and TMSD methylated standards (B) were resuspended in 25 mM ammonium acetate in chloroform:MeOH (1:1, v/v) and measured by nESI-DIMS. Ammonium adducts of un-methylated and methylated DolP species ranging from C80 to C95 are indicated.

MS-based lipid quantification typically relies on internal lipid standards, i.e. lipid species not present in the biological samples, added before the extraction procedure. Ideally, these standards resemble the analytes in their physico-chemical properties and behave similarly in derivatization, extraction, chromatographic behavior, and MS analysis. Isotopically labeled counterparts to endogenous lipids are the molecules of choice because standards can be used that are very close to the m/z range of the target lipid species. Since such standards are not commercially available for DolPs, we tested analytes that closely resemble DolPs but with m/z values that do not overlap those of the endogenous DolPs. Polyprenyl phosphates (PolPs) are structural analogs of DolP that are used in bacterial N-glycosylation and are also found in plants [2]. DolP and PolP share the phosphate head group structure and show a similar linear isoprene linkage, except for the α-isoprene unit, which is unsaturated in the case of PolPs. To establish the quantitative analysis of DolPs, we found the commercially available PolP C60 species a suitable standard since it can be used as an internal standard in a broad range of cellular matrices, including yeast, mammals, and archaea [46].

We performed the TMSD-based methylation analysis of PolP C60 as described for DolP. As observed for DolP, methylation of PolP resulted in a complete shift towards di-methylated PolP species (Fig. S3), and methylated PolP showed RPLC-MS properties similar to methylated DolPs. The EIC for the [M+NH_4_]^+^ ion of methylated PolP C60 showed a chromatographic elution at RT ~19.37 min (Fig. S4A), and the characteristic qualifier ion, assigned to a protonated adduct of the methylated phosphate with theoretical m/z 127.0155 (Fig. S4B).

To assess the TMSD methylation efficiency for DolPs, we compared the IS normalized peak intensities of un-methylated and TMSD-methylated C80, C85, C90, and C95 species in the DolP standard using nESI-DIMS measurements. Our analyses demonstrate that TMSD derivatization efficiency was ≥99% for all DolP species investigated, with a conversion to only di-methylated species under the given reaction conditions (Table S2). We found that DolP species were methylated to the same relative extent in the three independent replicates, as illustrated by the low relative standard deviation (RSD <1%). These results demonstrate that TMSD efficiently and selectively methylates the hydroxyl groups of phosphate residue of DolPs to di-methylated DolP species.

### Methylation of DolP improves chromatographic performance

The coupling of LC separation with MS detection offers better sensitivity, selectivity and peak capacity, and reduces the ion suppression observed in direct infusion nESI-DIMS [47]. In addition, LC-based separation of analytes reduces the complexity of biological lipid extracts and allows the enrichment of target analytes according to their physico-chemical properties. These advantages of LC-MS analysis are particularly important when it comes to the analysis of analytes present as minor components in a complex mixture, as is the case for DolP species in most biological samples. Among the LC separation methods commonly used for complex lipid mixtures, RPLC is the prevalent analytical technique [30]. RPLC is, however, unsuitable for the analysis of unmodified DolP species due to their wide range of lipophilicity (xlog P > 20) values. The methylation of phosphate groups of lipids via derivatization reduces the polarity of phospholipids, thus suggesting methylation to be a suitable method to enable RPLC analyses of DolPs. We, therefore, tested whether the methylation of DolP species makes these analytes amenable to RPLC separations, as has been previously shown for other anionic lipids [40, 48]. For the chromatographic separation of individual methylated DolP species we used a conventional C18 column prior to MS detection. We found that methylation of DolPs markedly improved the ability of their RP chromatographic retention and resolution and resulted in a narrower peak shape of methylated DolPs compared to their unmodified counterparts (Fig. S5). Importantly, no carryover was observed for methylated DolPs in the RPLC system. The improved chromatographic elution characteristics observed for the methylated DolP species is comparable to what was described for the analysis of methylated phospholipids using supercritical fluid chromatography-MS [44]. TMSD methylation combined with RPLC-MS analysis was also shown to be a powerful approach for detecting and quantifying phosphatidylinositol-(3,4,5)-trisphosphate species, which, comparable to DolPs, are present in cellular extracts only in very small amounts [35]. After methylation, we detected the DolP species C70-C105 species (Table 1, see Table S3,4 for species distributions) in the DolP standard by RPLC-MS analysis. A C65 species listed by the manufacturer was not detectable in the standard. A list of all annotated methylated DolP species characterized in this study, their theoretical and observed m/z values, molecular formulae and retention times are provided in Table 1.

An overlay of individually extracted ion chromatograms (EICs) of various DolP species obtained after RPLC-MS analysis of the methylated DolP standard demonstrates baseline resolution of DolP species with differing chain lengths within a retention time (RT) range of ~20-25 min (Fig. 2). Increasing isoprene units (with increased degree of unsaturation and carbon length) resulted in longer RTs of methylated DolP species (Table 1). This increase of RTs is attributed to an increase in the hydrophobic interactions of DolPs species with the stationary phase [49]. Notably, the isotopic peaks of the [M+NH_4_]^+^ adducts of DolP species were fully resolved at Rm/z200= 7×10^4^. For DolP C90, C95, C100, and C105 species, the M+1 isotope peak was more abundant than the monoisotopic M+0 peak (see Fig. S2 for DolP C90 and DolP C95), as is expected due to the increased contribution of ^13^C isotopes.

**Figure 2.**
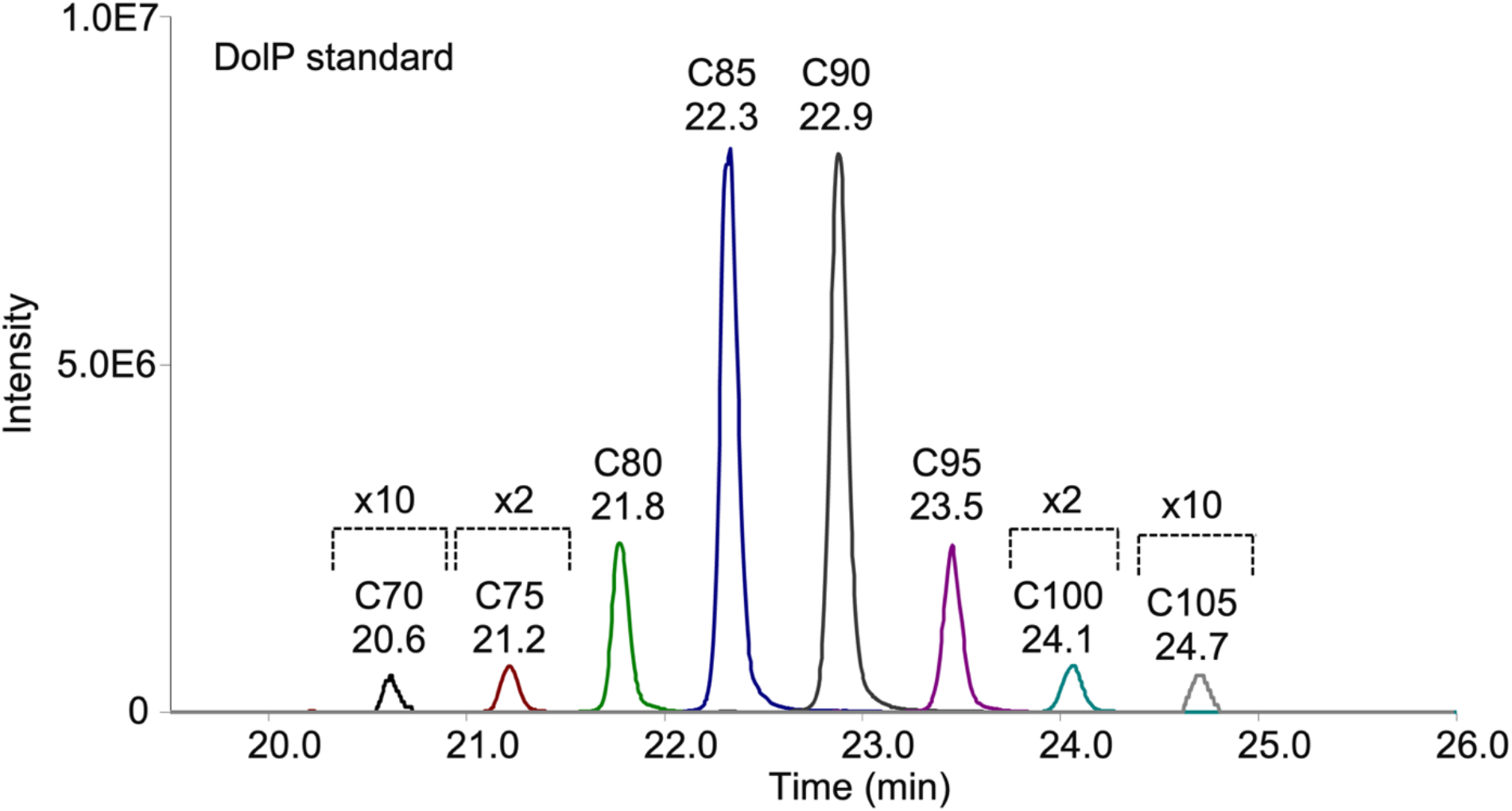
Separation and detection of methylated DolP species in DolP standard by RPLC-MS analysis. The DolP standard was subjected to methylation and RPLC-MS analysis. A representative overlay of EICs of di-methylated [M+NH_4_]^+^ ions of DolP species in standard is shown. The peaks are labeled with the retention times and correspond to the relevant EIC peaks for the specified DolP species. Low intensities were increased 2 or 10 times for visualization as indicated.

Having demonstrated the baseline separation of DolP species by MS analysis of [M+NH_4_]^+^ ions, we performed MS/MS analyses of the individual DolP species for further characterization, as exemplified for the DolP species C95 present in the DolP standard (Fig. 3). Following RPLC separation and MS detection, the EICs of DolP C95, detected as [M+NH_4_]^+^ and [M+H]^+^ ions, illustrate that the abundance of the [M+H]^+^ ion was typically ~30 fold lower than the corresponding [M+NH_4_]^+^ ion (Fig. 3A). [M+NH_4_]^+^ adducts were therefore used for the mass spectrometric characterization of DolPs.

**Figure 3.**
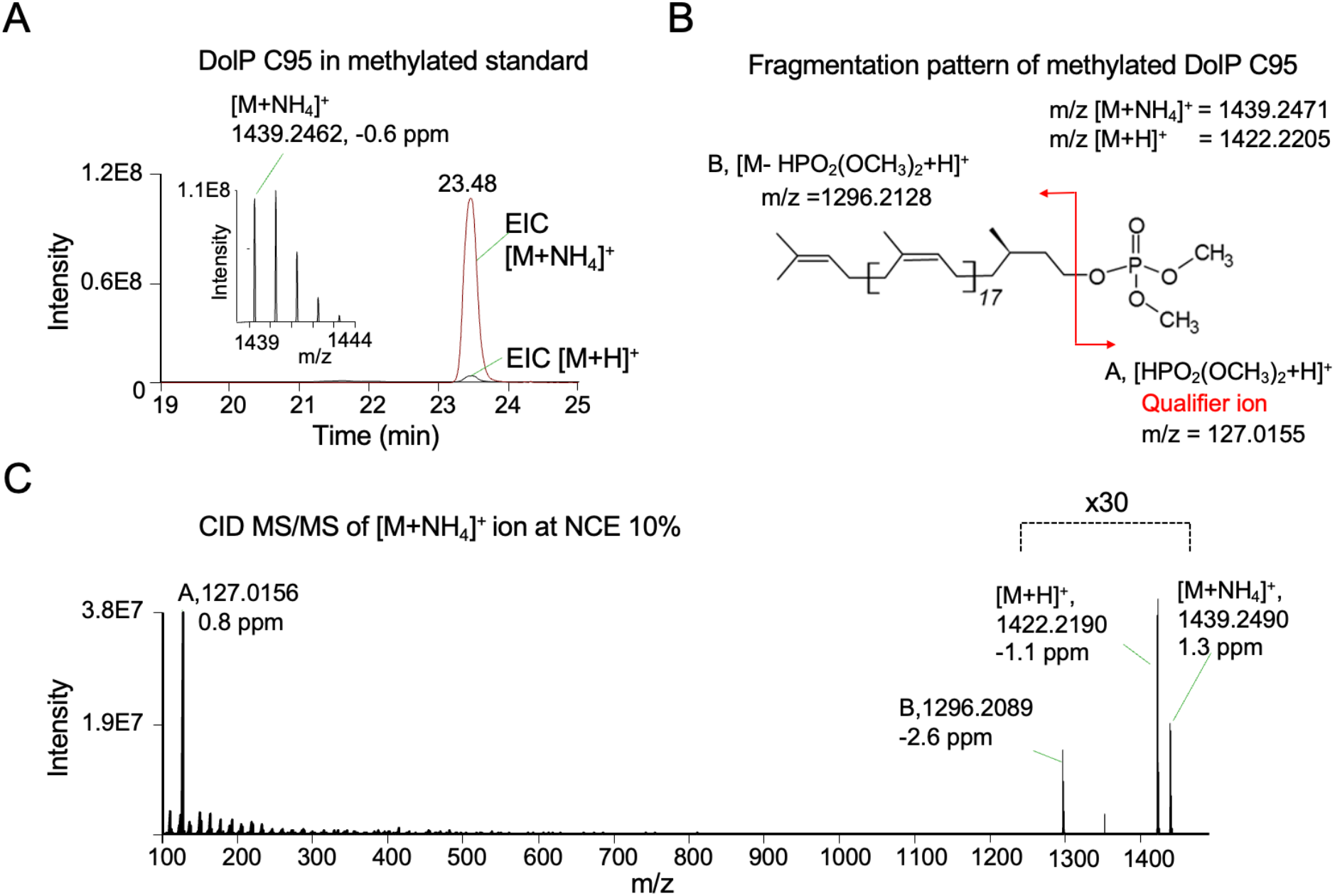
RPLC-MS characteristics of di-methylated DolP C95 species in DolP standard. (A) Overlay of EICs (with a narrow extraction window of ±5 ppm) of the [M+NH_4_]^+^ and [M+H]^+^ ions, with the corresponding MS spectrum of the ammonium adduct. (B) Fragmentation pattern of di-methylated DolP C95 as [M+NH_4_]^+^ ion. (C) MS/MS spectrum resulting from low collision energy (NCE 10%) induced dissociation of [M+NH_4_]^+^ showing the intact [M+NH_4_]^+^ and [M+H]^+^ ions and the characteristic qualifier ion. Capital letters A and B refer to the illustration of the fragmentation pattern shown in (B).

Collision energy (CE) induced dissociation of the precursor [M+NH_4_]^+^ ion using low CE (NCE 10%) resulted in the appearance of a highly abundant head group fragment ion in the MS/MS spectrum, corresponding to a protonated dimethylphosphate [HPO_2_(OCH_3_)_2_+H]^+^ ion (Fig. 3B,C), suggesting phosphodiester bond cleavage as the major fragmentation pathway, which was also previously demonstrated for sphingosine-1-phosphates with di-methylated phosphate headgroups [50]. This headgroup fragment ion was found in all MS/MS spectra of methylated DolP species, independent of the number of isoprene units, and was therefore used as a qualifier ion. In addition to the major fragment ion, other less abundant ions corresponding to the loss of ammonia and subsequent loss of a dimethyl phosphate ion were also observed (Fig. 3C).

The DolP species present in the DolP standard have different degrees of hydrocarbon chain unsaturation (13-21 double bonds per lipid) and range in length from 67 to 107 C-atoms. Their different physico-chemical properties (such as polarity or pKa) could consequently influence the derivatization and RP chromatographic behavior of DolPs. We, therefore, determined the relative distribution of the major DolP species C80, C85, C90 and C95, present in the DolP standard, comparing the profiles before and after methylation, and with or without RPLC separation prior to MS detection. Our data demonstrate that the DolP species profile is independent of the methylation reaction, and similar results are obtained for methylated DolPs in the presence or absence of RPLC separation (Table S3).

### Method validation

The optimized RPLC-MS method was validated according to ICH guidelines (see Supplementary Information for experimental details and results). First, we tested the RPLC-MS method for limits of quantification (LOQs), and determined the limit of detection (LOD) and the linearity range (Table S4, Fig. S6). We found that the limit of detection (LOD) for all DolP species was ≤2 pg on the column. Our method showed excellent linearity over an extensive range with correlation coefficient (r^2^) values of 0.99-1.00 for all DolP species. The linear range 0.05 to 800 pg/μL was sufficiently low to cover the quantification of endogenous DolP species at the cellular level. The LOQ of ≤10 pg (on column) is, to our knowledge, the most sensitive method for quantitative analysis of DolP species reported to date. Further, the method performance was assessed at three quality control (QC) levels, low (5), medium (20), and high (50) concentrations (in pg/μL) of methylated DolP standard spiked into methylated HeLa cells (matrix blanks). The specificity and selectivity of the method was tested by comparing matrix blanks (HeLa cells extracts) with and without methylation, and by comparing matrix blanks with and without spiked in methylated DolP/PolP C60 standards. For extraction recovery, QCs were spiked in blank solvent. The intra- and inter-day precision expressed as % CV (n = 3) was in the range of 1.1 - 7.9 %, and 0.5 - 11.2 %, respectively, with intra- and inter-day accuracy ranging from 100.1% - 110.4%, and 101.2% −110.3%, respectively (Table S5). For all DolP species, an extraction efficiency of ≥95% was found, except for DolP C105, which was only recovered in the high concentration sample (Table S6). Post-preparative autosampler stability results are summarized in Table S6 and indicate that there was no significant degradation of DolPs in these samples in the autosampler at 4 °C for over 20 h (Table S6). No background interference for methylated ions was found. Overall, the validated method is accurate, precise and acceptable for the analysis of biological samples. Taken together, TMSD methylation of DolPs prior to RPLC-MS offers several advantages such as the simple pre-column derivatization procedure, improved stability of derivatized samples, and enhanced sensitivity of detection by masking phosphates and improving ionization.

### DolP species profiling in biological samples

To determine the DolP content of HeLa cells, ~1×10^6^ cells were subjected to alkaline hydrolysis, extracted, methylated, and subjected to RPLC-MS analysis. The alkaline hydrolysis step is essential to ensure optimal recovery of DolP from biological samples. At the same time, oligosaccharides are released from dolichyl carriers and converted to DolPs [51]. Cellular DolP species were annotated by similarity matching of precursor m/z, RPLC retention times, presence of the qualifier head group fragment in the MS/MS spectrum, and similarity matching of isotope patterns of endogenous DolP species and standards. A representative RPLC-MS chromatogram, MS, and MS/MS analysis of endogenous DolP C95 from methylated HeLa cells extracts are shown in Fig. S7. In total, we detected eight DolP species in HeLa extracts, ranging from C70 to C105 (Fig. 4A), which were baseline-separated in their EIC profiles (Fig. S8A). DolP C95 was the most abundant DolP species at 54.1±0.2% of the total DolP pool, which is in agreement with earlier reports [46, 52, 53]. Due to the high sensitivity of our method, we could also identify DolP C70, C75 and C80 as minor species for the first time, accounting for 0.4±0.1%, 0.5±1% and 0.7±0.1% of total DolP species respectively. Normalizing DolP to PC, which contributes ~ 33% to membrane lipids in HeLa cells [54], revealed that the DolP content of HeLa membranes is in the range of 0.008% of membrane lipids and is thus present only in catalytic amounts.

**Figure 4.**
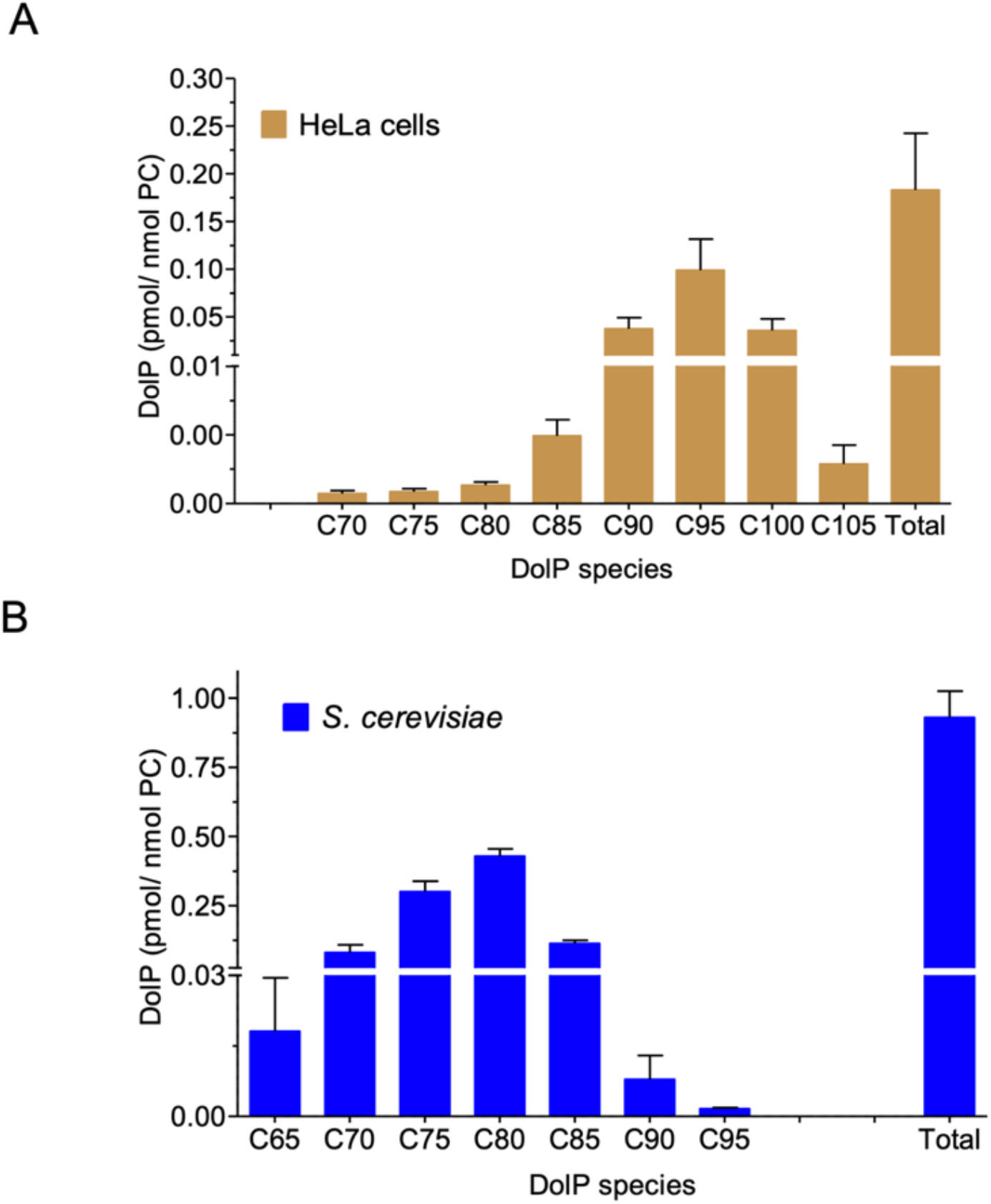
DolP species profiles determined in cellular extracts. HeLa (A) and *S. cerevisiae* (B) cells were subjected to alkaline hydrolysis and lipid extraction, methylation, and RPLC-MS analysis. DolP amounts were normalized to phosphatidylcholine (PC) as bulk membrane lipid. The inset on each graph shows minor lipid species. Data are presented as mean ± SD of three biological replicates.

Similarly, we characterized the DolP profile of *S. cerevisiae* cells. In contrast to HeLa cells, DolP species in *S. cerevisiae* range from DolP C65 to DolP C95, with DolP C75 and DolP C80 being the dominant lipid species (Fig. 4B, Fig. S8B). DolP C70 to C85 species together formed 31.7±0.9% and 45.5±2.9% of total DolPs, respectively, and their distribution mirrors that of the DolP precursor dolichol [55].

### Monitoring DolPs in SRD5A3-CDG patient fibroblasts

Having validated the DolP quantification in cellular extracts, we determined DolP levels in fibroblasts from patients suffering from a mutation in the steroid 5-*a*-reductase type 3 (SRD5A3) gene [55]. The gene encodes polyprenol reductase, which catalyzes the reduction of the alpha-isoprene of polyprenol in the final step of dolichol synthesis (Fig. S1). SRD5A3-CDG is associated with severe visual impairment, variable ocular anomalies (e.g. iris and optic nerve coloboma, congenital cataract, glaucoma), intellectual disability, cerebellar abnormalities, hypotonia, ataxia, ichthyosiform skin lesions, kyphosis, congenital heart defects, hypertrichosis and abnormal coagulation, affecting children and adults (for examples see [55–60]) and reflecting the central role of dolichol in cellular glycosylation. Fibroblasts from a SRD5A3-CDG patient presenting a severe phenotype were analyzed for their DolP content. Compared to healthy control fibroblasts, the DolP content was significantly decreased (41.5%) in the patient’s fibroblasts (Fig. 5A), supporting a key role of SRD5A3 in maintaining cellular DolP levels. The presence of a substantial residual amount of DolP in the patient’s fibroblasts is in line with a recent report suggesting a yet unidentified alternative pathway of dolichol synthesis [61].

**Figure 5.**
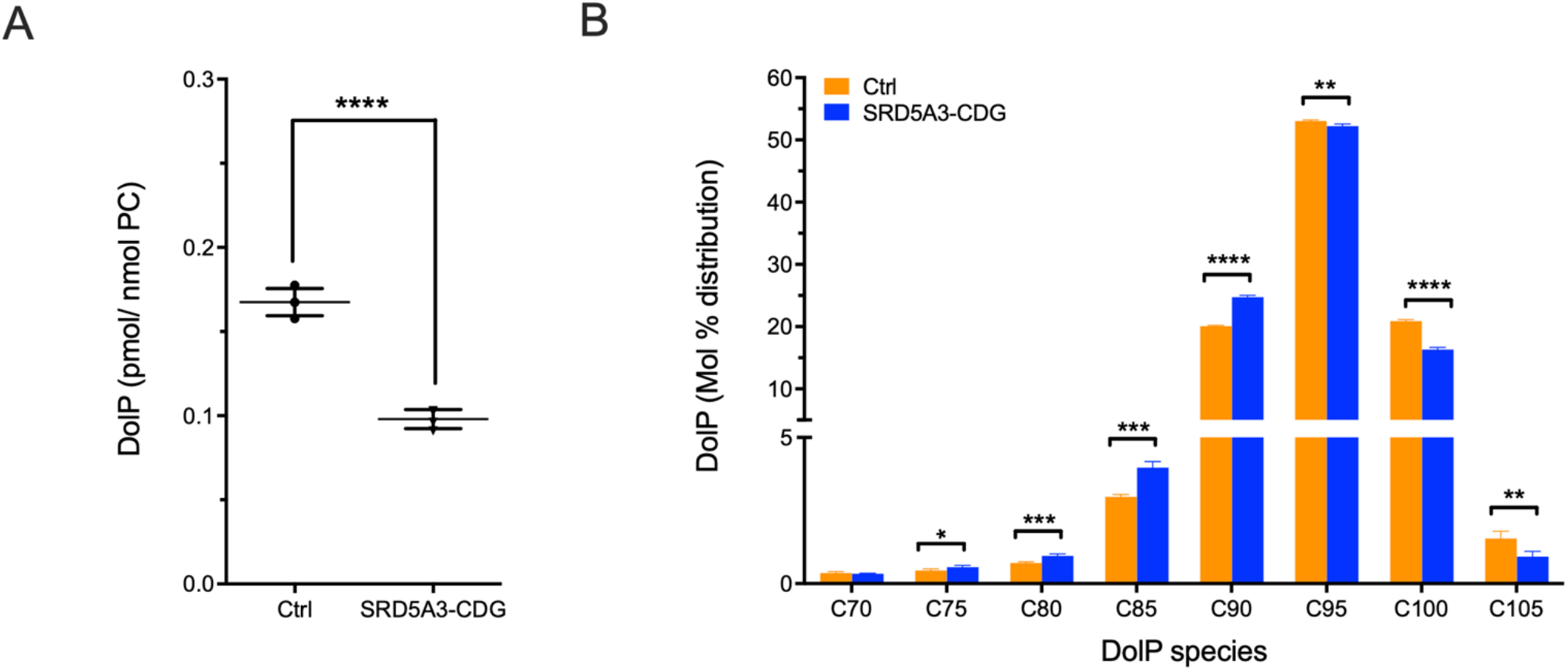
DolP homeostasis is perturbed in SRD5A3-CDG fibroblasts. Quantification of DolP content in SRD5A3-CDG patient fibroblasts. Patient fibroblasts (SRD5A3-CDG) and healthy control cells (Ctrl) were subjected to lipid extraction in the presence of internal standard, methylation, and RPLC-MS analysis. (A) Total DolP amounts were normalized to PC. (B) Distribution of DolPs ranging from C70 to C105 displayed as a percentage of all DolP species. Data are presented as mean ± SD of four biological replicates.

We also observed a shift towards the shorter DolP C75 - DolP C90 species, mainly at the expense of DolP C100 and DolP C105 (Fig. 5B). Inhibition of the DolP synthesis pathway by the small molecular drug lovastatin also led to a decrease in DolP species chain length in liver tissue of lovastatin treated mice [62] and in hepatocellular carcinoma samples in comparison to healthy human liver [63]. In all cases, the underlying molecular mechanisms are unclear and illustrate the importance of including DolP species profiling when studying DolP homeostasis in health and disease.

## Conclusions

Here we describe a method for rapid and highly sensitive quantitative analysis of DolP species in biological samples. DolP profilings are not yet implemented in comprehensive lipidomics analyses, mainly due to their very low abundance. By combining phosphate methylation with RPLC-MS separation, we established a workflow to quantify DolP species from as few as 10^6^ HeLa cells. Including the TMSD-based methylation allowed the quantification of DolP species over a broad range of lipophilicities. Due to its unprecedented sensitivity, our method now offers comprehensive coverage of DolP species ranging from C70 to C105, including minor species that were not detected in previous studies. Sample preparation has been simplified by implementing a rapid 3-step procedure, including hydrolysis, extraction, and methylation. The current method provides access to RPLC-MS analysis that was not readily accessible via the previous method. Of note, this method is also suitable for analyzing PolP species. The implementation of DolP species analysis into standard lipidomics workflows will help to discover alterations in DolP steady state levels, which are intricately linked with a number of lipophilic metabolites, including cholesterol and ubiquinone, via the mevalonate pathway thus connecting glycosylation processes with membrane homeostasis and cellular energy states. This approach also presents a promising diagnostic and/or therapeutic application for pathophysiologies such as Alzheimer’s disease that are associated with altered DolP levels in body fluids, including urine, plasma or liquor.

## Supporting Information

The Supplementary Information contains Supplementary Methods, Supplementary Figures, and Tables. All lipidomics data have been deposited to the Metabolights database, where they are available under accession number MTBLS4095 for RPLC-MS data, and MTBLS4228 for nESI-DIMS data.

## Author Contributions

D.K. and B.B.: Conceptualization, methodology, investigation, Writing: original draft preparation, review, editing. D.K.: Formal analysis, data curation, visualization. B.B.: Supervision, funding acquisition. P.P.: Writing: Review and editing. F.K., L.B., and C.T.: Cell culture preparation of biological samples. C.T.: Funding acquisition. All authors contributed to and approved the manuscript.

## Acknowledgments

This project was funded by the Deutsche Forschungsgemeinschaft (DFG, German Research Foundation) – Project-ID 289991887 – Forschungsgruppe FOR 2509, project 02 and – Project-ID 112927078 – Transregio TRR83, project 01). We thank Timo Sachsenheimer for PC measurements, Iris Leibrecht and Alexia Herrmann for technical assistance, and Ingo Amm for providing *S. cerevisiae* cells.

