## Supplemental Data for "Simple, rapid, and sensitive quantification of dolichyl phosphates using phosphate methylation and reverse-phase liquid chromatography-high resolution mass spectrometry"

Kale et al.

| Table of contents | |
| --- | --- |
| Additional experimental information and results on validating the novel approach for MS-based DolP analysis. | |
| Figure S1 | DolP synthesis in mammals. DolP synthesis in mammals is mainly localized to the cytoplasmic side of the ER. The biosynthesis of DolP starts with acetyl-CoA and proceeds via the mevalonate pathway. Farnesyl diphosphate serves as a branching point for the synthesis of DolP, cholesterol ubiquinone and heme and is also used for protein prenylation. Drugs such as statins to target cholesterol biosynthesis consequently also impact DolP synthesis and downstream glycosylation processes. Polyprenol is reduced to DolP by Steroid 5-α-reductase 3 (SRD5A3). The defect at this stage results in SRD5A3-CDG (MIM# 612379). |
| Figure S2 | Representative narrow m/z range positive ion nESI-DIMS spectra of C80-C95 species in DolP standard before (A) and after methylation (B). For theoretical m/z values of ammonium adducts of these un-, mono-, di-methylated DolP ions please see Table S1. |
| Figure S3 | Positive ion narrow m/z range nESI-DIMS spectra of PolP C60 standard after methylation. |
| Figure S4 | RPLC-MS characteristics of di-methylated PolP C60 standard. A, EIC (±5 ppm) of the [M+NH_4_]^+^ ion, with corresponding MS spectrum. Theoretical fragmentation pattern of the di-methylated PolP C60. B, MS/MS spectrum resulting from low collision energy (NCE 10%) induced dissociation of [M+NH_4_]^+^ ion, showing the characteristic qualifier/head group fragment ion. Capital letters A and B refer to the illustration of the fragmentation pattern shown in (A). |
| Figure S5 | Comparison of EICs of [M+NH_4_] ^+^ ion for RPLC-MS chromatographic performance of C80-95 species in DolP standard before and after methylation |
| Figure S6 | Linearity curves for the RPLC-MS analysis of C70-105 species (on column) in DolP standard. |
| Figure S7 | RPLC-MS characteristics of endogenous di-methylated DolP C95 from HeLa cells. A, EICs (±5 ppm) of the [M+NH_4_]^+^ ions, with corresponding MS spectrum. B, MS/MS spectrum resulting from low collision energy (NCE 10%) induced dissociation of [M+NH_4_]^+^ ion, with fragmentation pattern showing the characteristic qualifier ion and other fragment ions. |
| Figure S8 | EICs of DolP species profiled using RPLC-MS analysis in cellular extracts of (A) HeLa cells and (B) *S. cerevisiae* |
| Table S1 | Theoretical m/z list of un-methylated (+0 Me), mono-methylated (+1 Me), di-methylated (+2 Me) ions of C80-C95 species in DolP standard |
| Table S2 | % Methylation efficiency of C80-C95 species in DolP standard measured using nESI-DIMS. |
| Table S3 | % Distribution of C80-C95 species in DolP standard with and without RPLC separation and/or methylation, followed by MS detection. |
| Table S4 | % Composition of C70-C105 species in methylated DolP standard, calibration curve parameters, instrumental limit of detection (LOD) of the RPLC-MS method for the simultaneous determination of targeted DolP species. |
| Table S5 | Results from intra- and inter-day accuracy and precision of RPLC-MS analysis of DolP standard in HeLa cells extracts |
| Table S6 | Results from determination of extraction recovery (n=3) in blank solvent and post-preparation stability of RPLC-MS analysis of DolP standard in HeLa cells extracts in autosampler for 20h (n=3). |

### Experimental Section

### Method validation

The methylated DolP standard was reconstituted in methanol, and a dilution series was prepared from 0.05 to 800 pg/µL. For each DolP species, the linearity was assessed by plotting the analyte-to-internal standard peak area against the nominal concentration of individual species calculated based on their mol % distribution in the DolP standard (Table S4, Fig. S6). Calibration curves (n=3) consisting of four to seven calibrator levels were constructed by simple linear regression analysis. For each DolP species, the lower limit of detection (LOD) was chosen as the calibration concentration when a signal-to-noise ratio is at least 5:1.

Validation of the method was assessed at three-level quality control (QC), at low (50), medium (200), and high (500) pg/µL of methylated DolP standard mixture. The intra- and inter-day assays were determined by analyzing the prepared samples on the same day and on three consecutive days. For precision and accuracy evaluations, samples were prepared by spiking the QC levels to the methylated Hela cells extracts (n=3). Accuracy is assessed as an agreement between the back-calculated concentration of individual analyte species in the spiked samples and their nominal concentrations, whereas precision is expressed as the coefficient of variation (%CV) for both intra-day and inter-day analysis values are shown in Table S3. For DolP species extraction recovery (Table S6), samples were prepared as pre-spikes and post-spikes at high and low QC levels in a pure solvent. The samples were extracted as described in the main text experimental section. The post-preparative stability (Table S6) was determined at low and high QC levels. These QC levels were spiked to methylated HeLa cell extracts, kept at 4 ºC for 20 h in an autosampler. The results were compared to conditions using freshly prepared samples with the same nominal concentration.

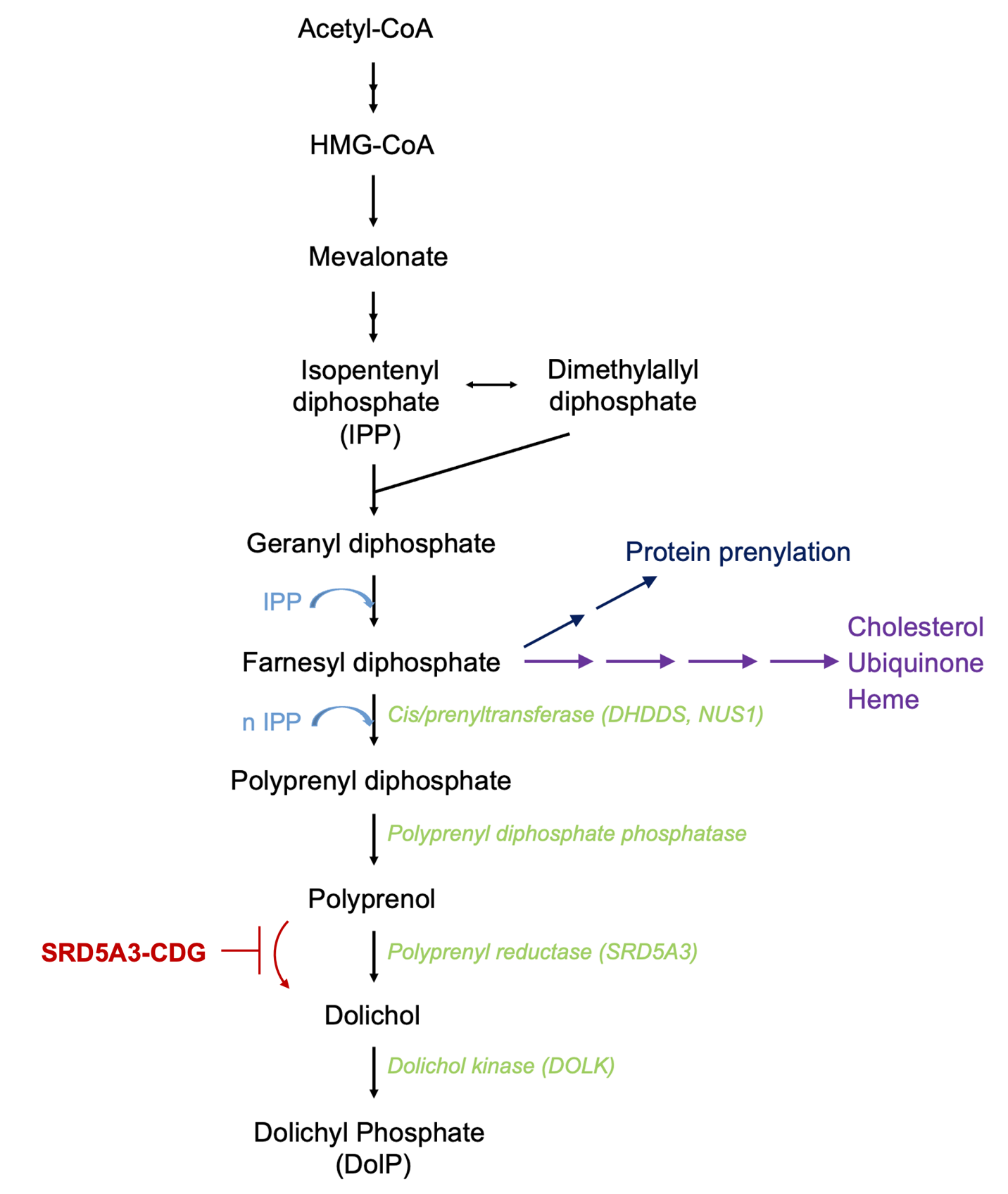

Figure S1

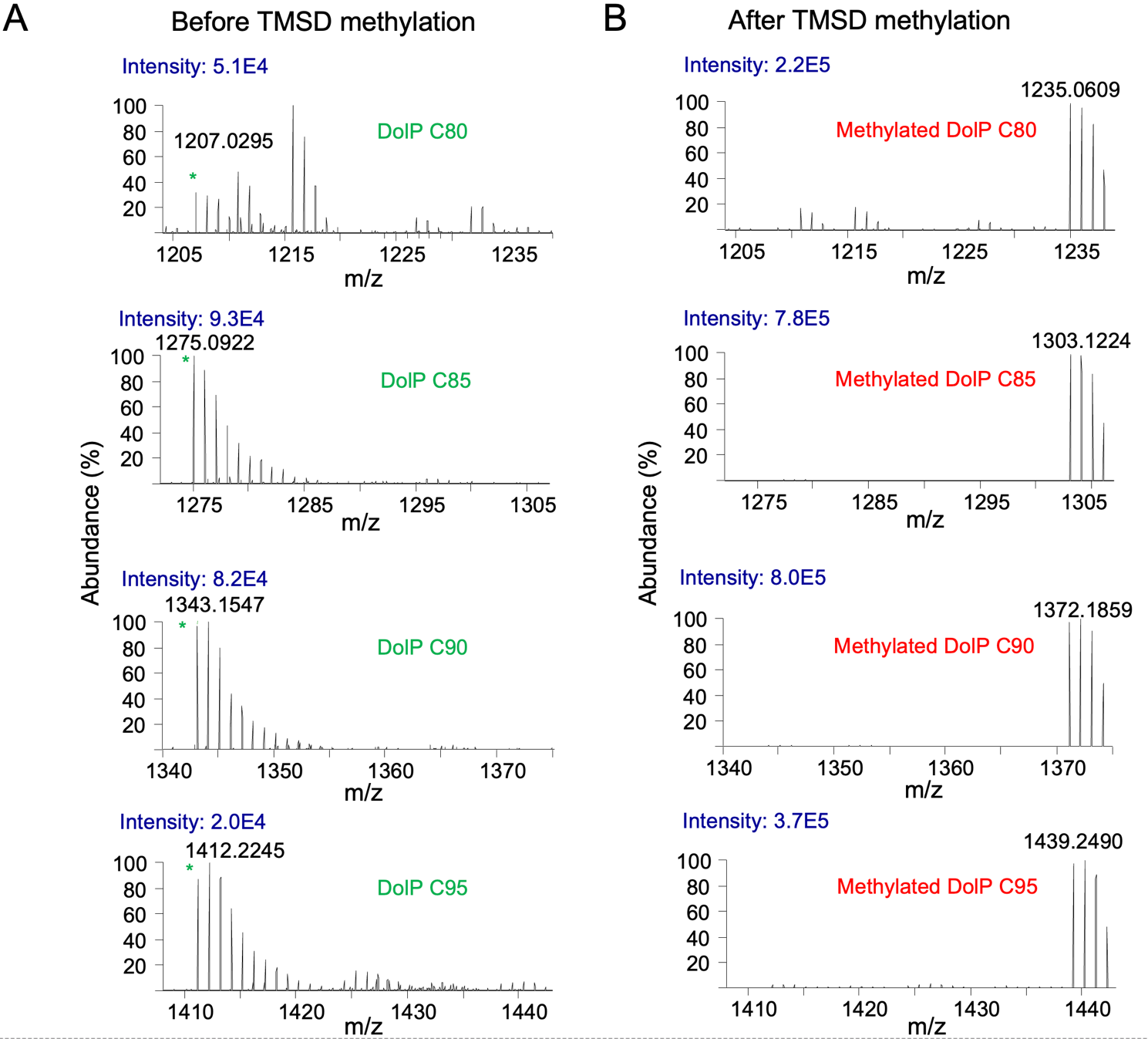

Figure S2

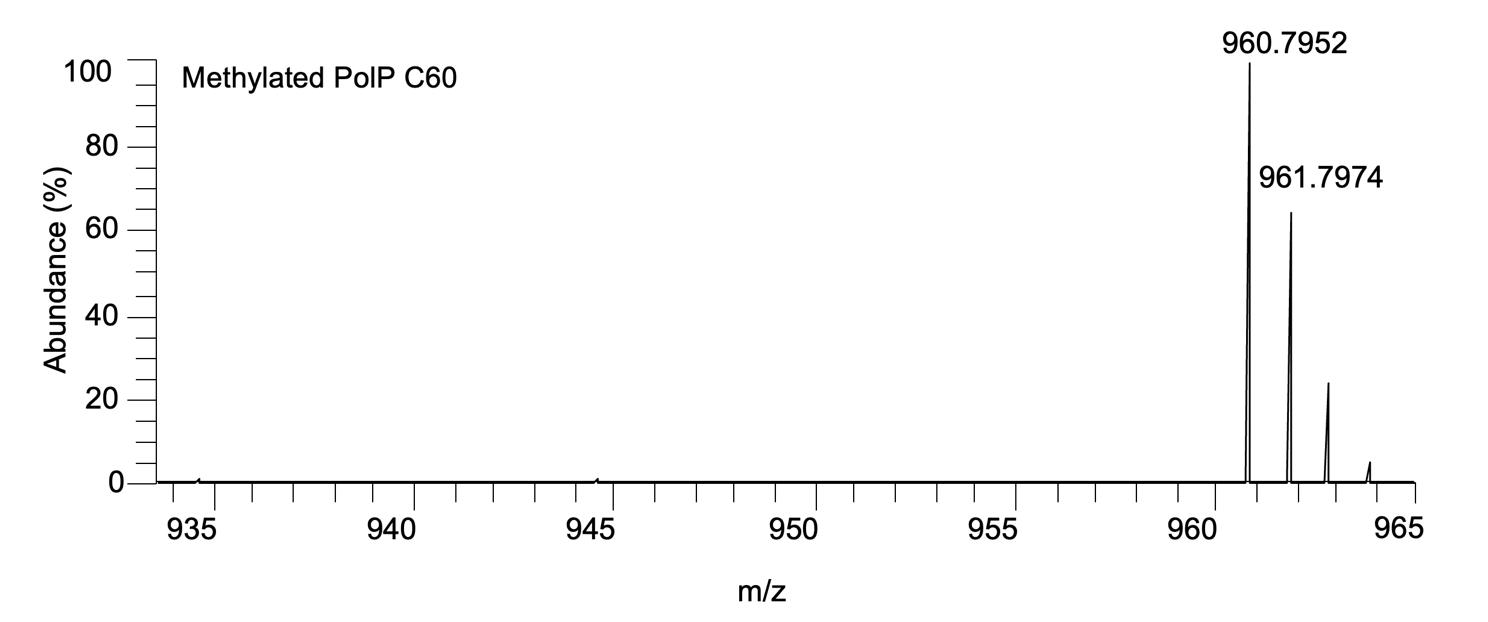

Figure S3

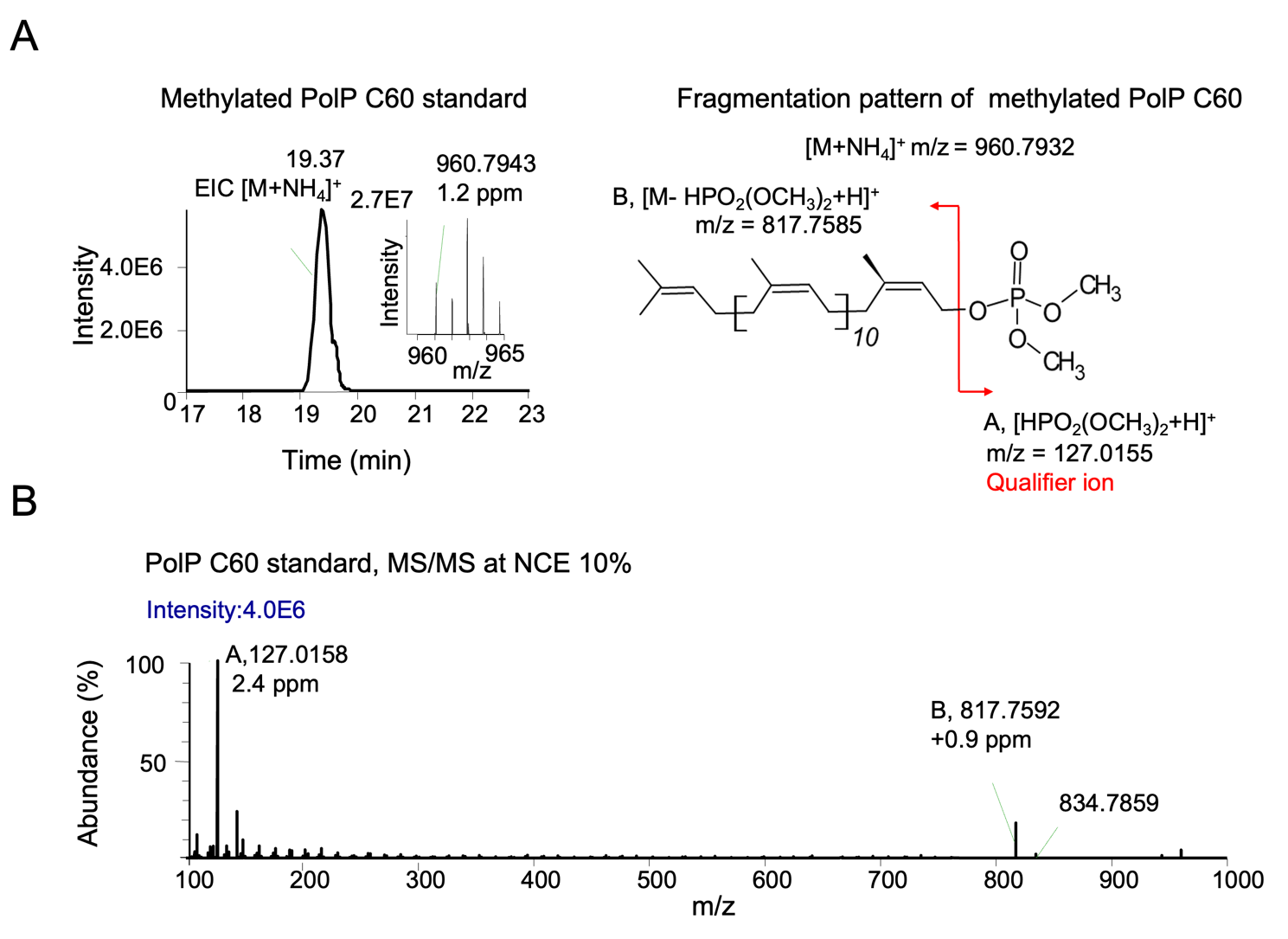

Figure S4

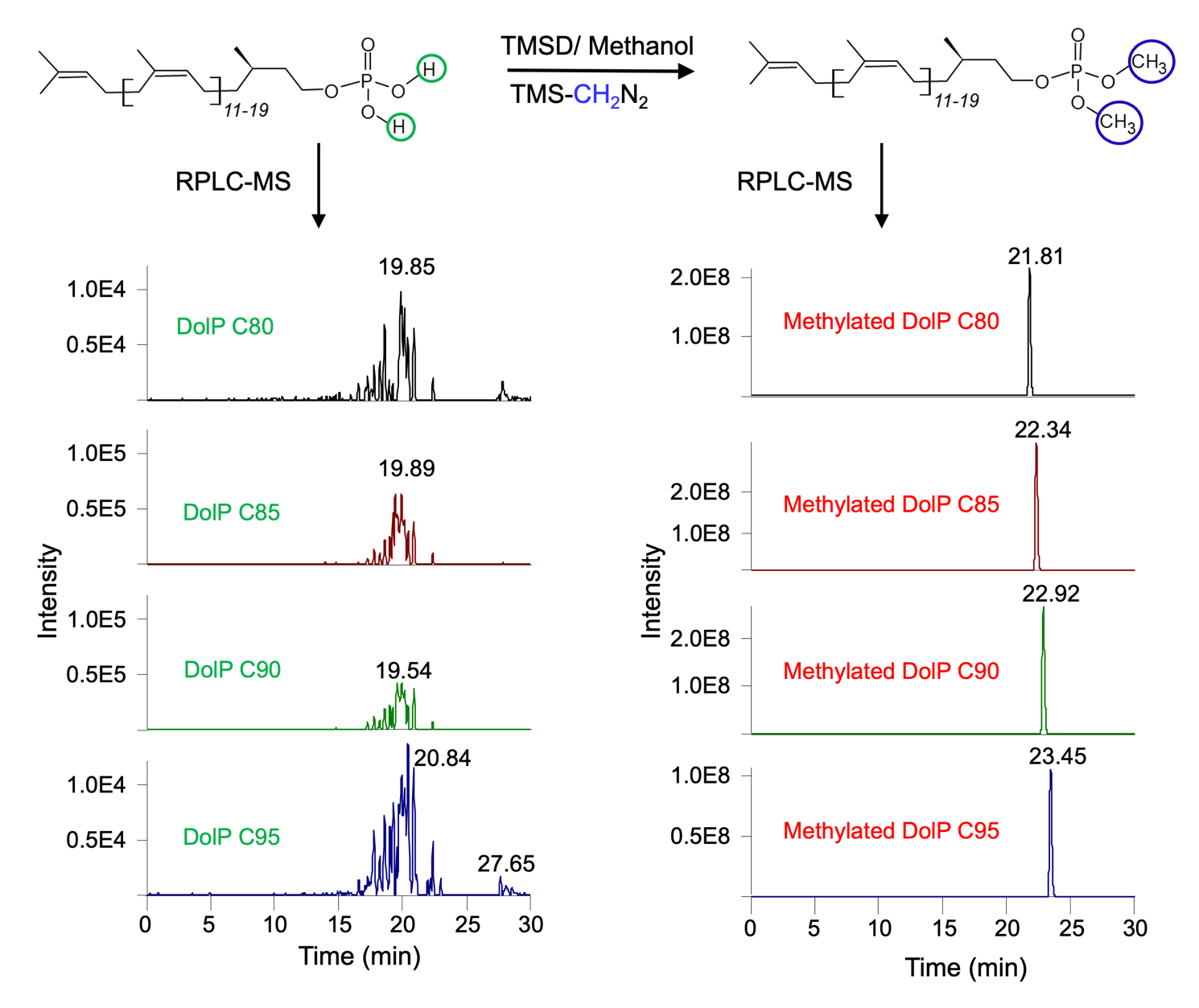

Figure S5

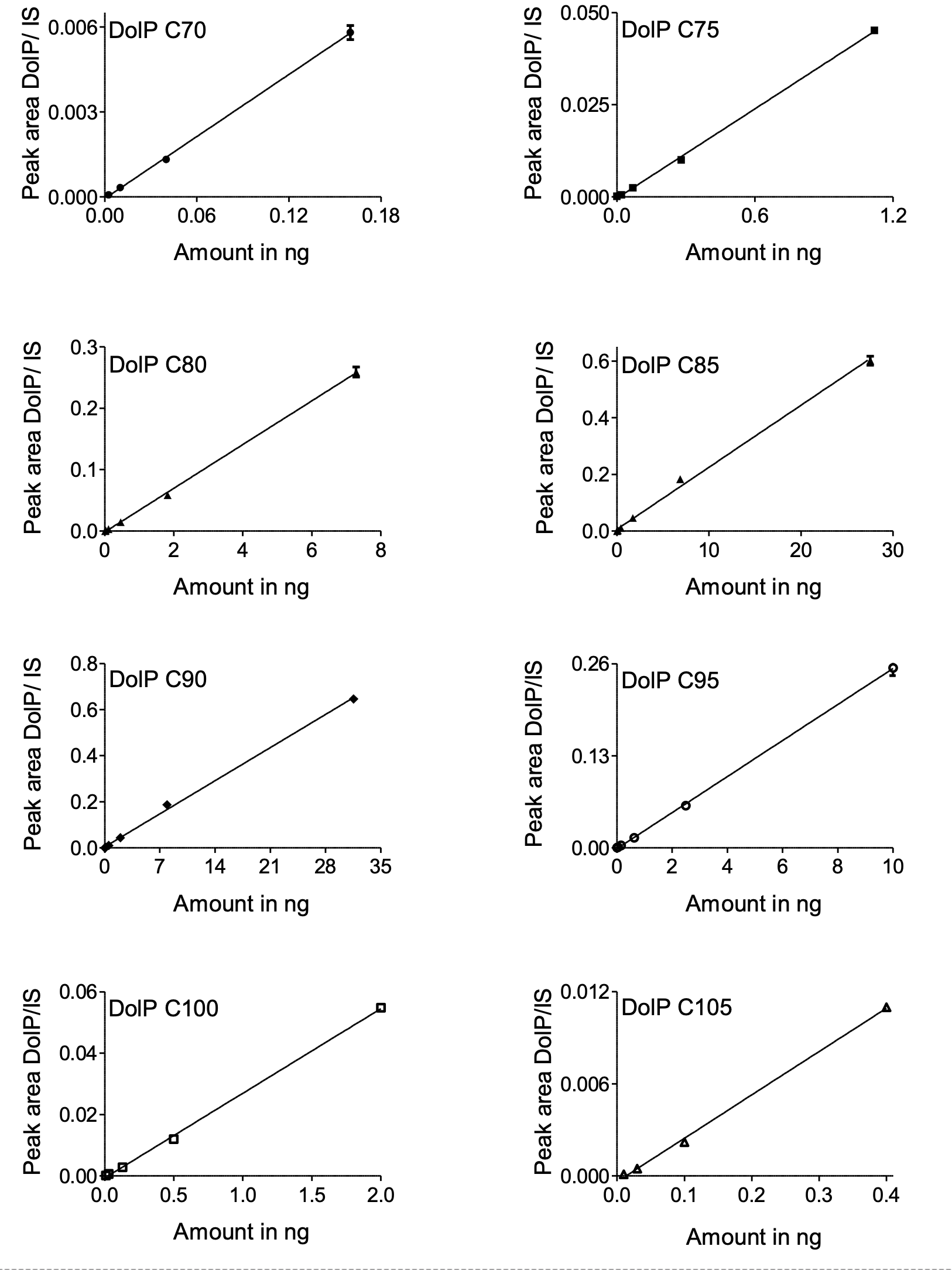
Figure S6

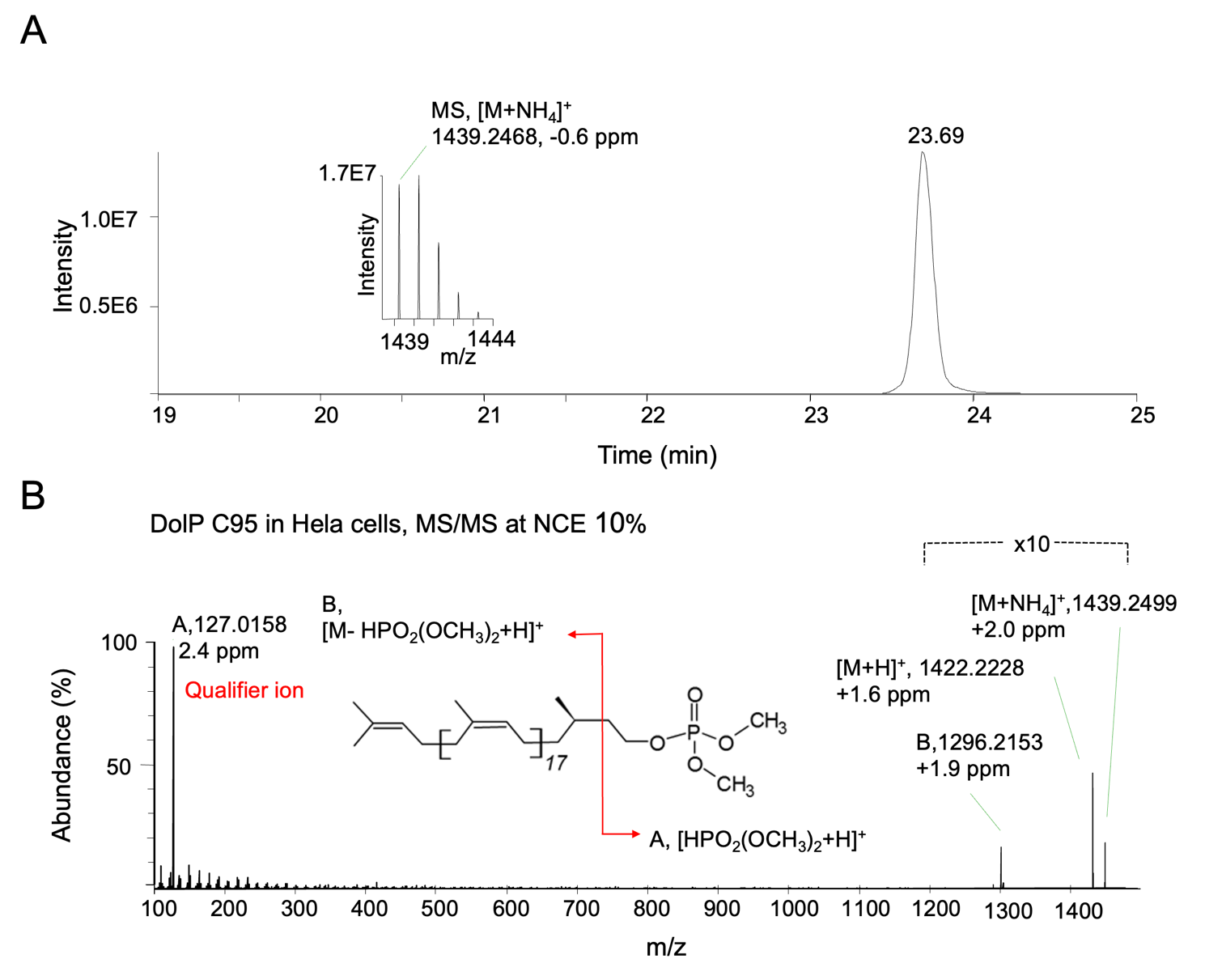

Figure S7

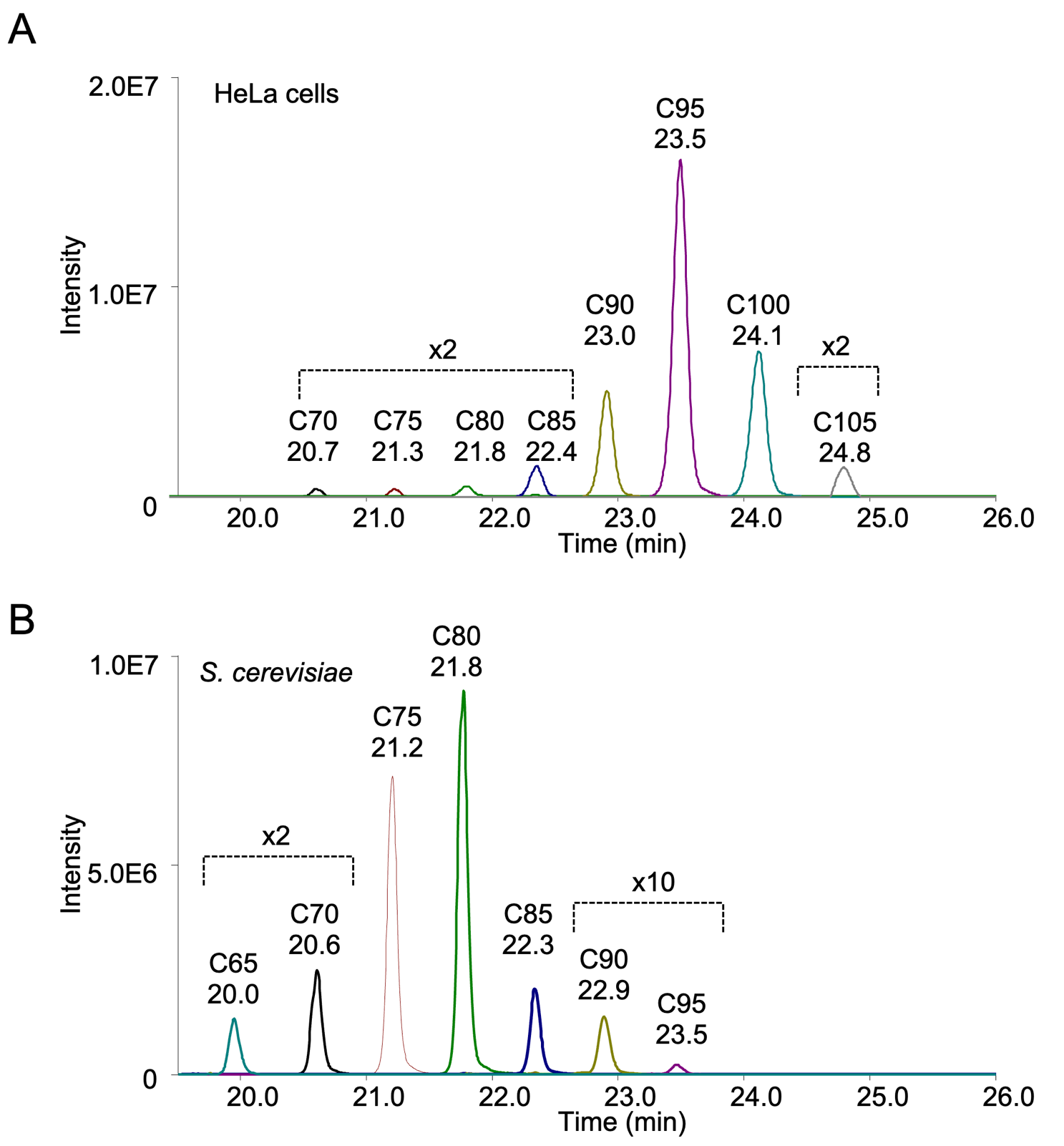

Figure S8

### Table S1

| DolP species  chain length | m/z list of species in DolP standard | | | | |
| --- | --- | --- | --- | --- | --- |
|  | +0 ^a^ Me | +1 ^a^ Me | | +2 ^a^ Me | |
| C80 | 1207.0280 | | 1221.0436 | | 1235.0593 |
| C85 | 1275.0906 | | 1289.1062 | | 1303.1219 |
| C90 | 1343.1532 | | 1357.1688 | | 1371.1845 |
| C95 | 1411.2158 | | 1425.2314 | | 1439.2471 |

^a^ The number of methylations in DolP

### Table S2

| DolP species  chain length | %^a^ of methylated compounds ^c^ | |
| --- | --- | --- |
|  | +0^b^Me | +2^b^Me |
| C80 | 0 ± 0.34 | 100 ± 0 |
| C85 | 0.82 ± 0.46 | 99.02 ± 0.16 |
| C90 | 0.33 ± 0.57 | 99.06 ± 0.2 |
| C95 | 0 ± 0 | 100 ± 0 |

^a^

$$\% of methylation efficiency=\left( 1-\frac{Intensity of intact DolP species after methylation}{Intensity of intact DolP species before methylation} \right)*100\%$$

^b^ Number of methylations in DolP

^c^ Values are mean ± SD, (n = 3)

### Table S3

| DolP species  chain length | % ^a^ Distribution of species in DolP standard | | |
| --- | --- | --- | --- |
|  | without methylation | with methylation | |
|  | nESI-DIMS | nESI-DIMS | ^b^ RPLC-MS |
| C80 | 11.1 ± 0.2 | 11.0 ± 0.3 | 10.4 ± 0.3 |
| C85 | 38.4 ± 0.3 | 36.8 ± 0.5 | 37.1 ± 1.5 |
| C90 | 38.3 ± 0.1 | 39.3 ± 0.3 | 40.4 ± 1.1 |
| C95 | 12.2 ± 0.3 | 12.9 ± 0.5 | 12.2 ± 0.3 |

^a^ Values represent peak areas of species, normalized to total peak area and are expressed as mean ± SD, n=3 independent methylations /non-methylation measurements.

^b^ For detailed distribution of methylated species on RPLC-MS refer to Table S4.

**Table S4**

| DolP species chain length | % Distribution ^a^ in DolP standard | Calibration range^a^ (pg on column) | LOD^a^ (pg on column) | r^2^ | Calibration Curve equation |
| --- | --- | --- | --- | --- | --- |
| C70 | 0.2 ± 0 ^b^ | 3 - 160 | 1 | 0.99 | *y =* 0.036*x -* 0.0001 |
| C75 | 1.4 ± 0.1 ^b^ | 4 - 1120 | 1 | 1.00 | *y =* 0.041*x -* 0.0004 |
| C80 | 9.1 ± 0.3 | 7 - 7280 | 2 | 1.00 | *y =* 0.036*x -* 0.0016 |
| C85 | 34.4 ± 1.4 | 7 - 27520 | 2 | 1.00 | *y =* 0.022*x +* 0.0060 |
| C90 | 39.4 ± 1.0 | 8 - 31520 | 2 | 1.00 | *y =* 0.021*x +* 0.0042 |
| C95 | 12.5 ± 0.3 | 10 - 10000 | 2 | 1.00 | *y =* 0.025*x -* 0.0012 |
| C100 | 2.5 ± 0.0 ^b^ | 8 - 2000 | 2 | 1.00 | *y =* 0.028*x -* 0.0006 |
| C105 | 0.5 ± 0.1 ^b^ | 6 - 400 | 2 | 1.00 | *y =* 0.028*x -* 0.0003 |

^a^ Amount of different species in a DolP standard, expressed as the ratio of the peak area of each DolP species to the total area of all DolP species, determined in three independent methylation experiments and represent mean ± SD

^b^ Additional DolP species detected in RPLC-MS but not in nESI-DIMS analyses

### Table S5

|  | Nominal concentration (pg/µL) | Accuracy (n=3) | | Precision (n=3) | |
| --- | --- | --- | --- | --- | --- |
| DolP species chain length |  | Inter-day | Intra-day | Inter-day | Intra-day |
|  |  | (Mean ± SD,%) | (Mean ± SD,%) | (CV,%) | (CV,%) |
| C70 | Low 50 | 107.2 ± 4.6 | 104.5 ± 4 | 3.78 | 4.28 |
|  | Middle 200 | 98.5 ± 3.7 | 96.1 ± 3 | 3.16 | 3.75 |
|  | High 500 | 102.7 ± 0.9 | 102.5 ± 1.1 | 1.06 | 0.87 |
| C75 | Low 50 | 105 ± 1 | 103.3 ± 2.9 | 2.84 | 0.91 |
|  | Middle 200 | 96.4 ± 4.4 | 96.6 ± 3.4 | 3.51 | 4.6 |
|  | High 500 | 100.1 ± 2.5 | 101.7 ± 1.9 | 1.9 | 2.52 |
| C80 | Low 50 | 99.2 ± 1.5 | 99.8 ± 2.8 | 2.81 | 1.5 |
|  | Middle 200 | 95 ± 5.2 | 96.6 ± 1.8 | 1.91 | 5.5 |
|  | High 500 | 99.5 ± 3.2 | 101.7 ± 2 | 1.94 | 3.18 |
| C85 | Low 50 | 99.8 ± 3.4 | 98.9 ± 3.7 | 3.77 | 3.39 |
|  | Middle 200 | 96.7 ± 5.9 | 97.3 ± 1.3 | 1.33 | 6.13 |
|  | High 500 | 100.5 ± 0.6 | 102.2 ± 2.2 | 2.15 | 0.63 |
| C90 | Low 50 | 100.3 ± 3 | 99.3 ± 4.1 | 4.1 | 2.98 |
|  | Middle 200 | 95.7 ± 4.4 | 97.4 ± 2.6 | 2.63 | 4.55 |
|  | High 500 | 100.1 ± 1.7 | 104.9 ± 4.3 | 4.1 | 1.68 |
| C95 | Low 50 | 104.5 ± 11.7 | 105.1 ± 4.1 | 3.86 | 11.15 |
|  | Middle 200 | 98.3 ± 4.4 | 101.2 ± 4 | 3.97 | 4.51 |
|  | High 500 | 98 ± 0.5 | 106.7 ± 7.6 | 7.13 | 0.46 |
| C100 | Low 50 | 102.2 ± 11.4 | 101.6 ± 4.8 | 4.77 | 11.19 |
|  | Middle 200 | 100.9 ± 4 | 103.5 ± 3.6 | 3.51 | 3.95 |
|  | High 500 | 99.6 ± 0.6 | 109.1 ± 8.3 | 7.62 | 0.61 |
| C105 | Low 50 | 110.4 ± 5.5 | 108.0 ± 7.5 | 6.71 | 7.66 |
|  | Middle 200 | 103.7 ± 3.6 | 106.2 ± 2.7 | 3.42 | 3.41 |
|  | High 500 | 100.7 ± 1.9 | 110.3 ± 8.3 | 7.88 | 1.63 |

### Table S6

|  | QC level | Extraction recovery (n=3) | | Stability (n=3) | |
| --- | --- | --- | --- | --- | --- |
| Analyte | (pg/µL) | Mean ± SD | RSD % | Mean ± SD | RSD % |
| C70 | Low (50) | 95.5 ± 1.8 | 1.9 | 119.6 ± 0.8 | 0.7 |
|  | High (500) | 106.4 ± 2.6 | 2.4 | 119.3 ± 1.6 | 1.3 |
| C75 | Low (50) | 103.6 ± 8.2 | 7.9 | 121.7 ± 1.4 | 1.2 |
|  | High (500) | 111.2 ± 4.5 | 4.1 | 118.5 ± 3.2 | 2.7 |
| C80 | Low (50) | 103.9 ± 8.9 | 8.5 | 120.5 ± 0.1 | 0.1 |
|  | High (500) | 115.3 ± 4.5 | 3.9 | 118.8 ± 2.3 | 2 |
| C85 | Low (50) | 112.2 ± 8.5 | 7.6 | 122.2 ± 1 | 0.8 |
|  | High (500) | 123.4 ± 6 | 4.8 | 124.9 ± 6.4 | 5.1 |
| C90 | Low (50) | 118 ± 9.1 | 7.7 | 123.8 ± 2 | 1.6 |
|  | High (500) | 125.6 ± 5.9 | 4.7 | 118.9 ± 7.3 | 6.1 |
| C95 | Low (50) | 123.3 ± 11.9 | 9.7 | 123.6 ± 3.2 | 2.6 |
|  | High (500) | 123.4 ± 6.2 | 5 | 113.8 ± 3.2 | 2.9 |
| C100 | Low (50) | 134.6 ± 12.8 | 9.5 | 122.3 ± 4.1 | 3.4 |
|  | High (500) | 121.6 ± 8 | 6.6 | 100.1 ± 2 | 2 |
| C105 | Low (50) | ND^a^ | 0 | 92.4 ± 15.9 | 17.2 |
|  | High (500) | 95.5 ± 1.8 | 1.9 | 99.9 ± 4.8 | 4.8 |

^a^ Not detected
